## Supplementary_file_1 for "Mutational biases promote neutral increases in the complexity of protein interaction networks following gene duplication"

Two molecules of A form the dimer AA. Concentrations are at a steady state determined by the law of mass action according to this reaction network :

| | | |
|---|---|---|
| =>sA | A | =>dA |
| 2A | r1<=>r2 | AA |
| | AA | =>dAA |

where $s_A$ is the synthesis rate of A (units of concentration*time^-1), $r_1$ is the dissociation rate of AA (units of concentration^-1*time^-1), $r_2$ is the association rate of A, and $d_A, d_{AA}$ are degradation rates (units of time^-1).

Let cA, cAA denote the concentrations of the two molecular species :

Because at equilibrium the concentration of A is constant :

**Equation 1 :** $s_A + 2 r_1 c_{AA} = d_A c_A + 2 r_2 c_A^2$

Because at equilibrium the concentration of AA is constant :

**Equation 2 :** $r_2 c_A^2 = d_{AA} c_{AA} + r_1 c_{AA}$

Starting from equation 2, we get an expression for $c_{AA}$ :

$$r_2 c_A^2 = d_{AA} c_{AA} + r_1 c_{AA}$$

$$0 = (d_{AA} + r_1) c_{AA} - r_2 c_A^2$$

$$c_{AA} = \frac{r_2 c_A^2}{d_{AA} + r_1}$$

With $k_{AA}$ defined as $\frac{r_2}{d_{AA} + r_1}$ :

$$c_{AA} = k_{AA} c_A^2$$

Starting from equation 1 :

$$s_A + 2 r_1 c_{AA} = d_A c_A + 2 r_2 c_A^2$$

$$0 = 2 r_2 c_A^2 + d_A c_A - s_A - 2 r_1 c_{AA}$$

By substituting $c_{AA}$:

$$0 = 2 r_2 c_A^2 + d_A c_A - s_A - 2 r_1 k_{AA} c_A^2$$

From the definition $k_{AA}=\frac{r_2}{d_{AA}+r_1}$ , we get $r_2-r_1 k_{AA}=d_{AA} k_{AA}$ and therefore :

$$0=2r_2 c_A^2+d_A c_A-s_A-2 r_1 c_A^2 k_{AA}$$

$$0=2r_2 c_A^2+d_A c_A-s_A-2 r_1 c_A^2 k_{AA}$$

$$0=2c_A^2\left(d_{AA} k_{AA}\right)+d_A c_A-s_A$$

Applying the quadratic formula :

$$c_A=\frac{-b\pm\sqrt{b^2-4ac}}{2a}=\frac{-d_A\pm\sqrt{d_A^2-4\left(2d_{AA}k_{AA}\right)\left(-s_A\right)}}{2\left(2d_{AA}\right)\left(k_{AA}\right)}$$

$$c_A=\frac{-d_A\pm\sqrt{d_A^2+8d_{AA}k_{AA}s_A}}{4d_{AA}k_{AA}}$$

In the formula above, the coefficients are such that $a>0$ , $b\geq 0$ , $c\leq 0$ and thus $b\leq\sqrt{b^2-4ac}$ . It follows that $-b+\sqrt{b^2-4ac}$ is non-negative, while $-b-\sqrt{b^2-4ac}$ is non-positive. We therefore choose $-b+\sqrt{b^2-4ac}$ to obtain a non-negative value of $c_B$ .

$$c_A=\frac{-d_A+\sqrt{d_A^2+8d_{AA}k_{AA}s_A}}{4d_{AA}k_{AA}}$$
