## Supplementary_file_2 for "Mutational biases promote neutral increases in the complexity of protein interaction networks following gene duplication"

Two molecules $A$ and $B$ form the dimers $AA$ , $AB$ and $BB$ . Concentrations are at a steady state determined by the law of mass action according to this reaction network:

=>sA **A** =>dA

=>sB **B** =>dB

**2A** r1<=>r2 **AA**

**2B** r3<=>r4 **BB**

**A + B** r5<=>r6 **AB**

**AA** =>dAA

**BB** =>dBB

**AB** =>dAB

where $s_A, s_B$ are synthesis rates (units of concentration * time^-1), $r_1, r_3, r_5$ are dissociation rates (units of time^-1), $r_2, r_4, r_6$ are association rates (units of concentration^-1 * time^-1), and $d_A, d_{AA}, d_{AB}, d_{BA}, d_{BB}, d_B$ are degradation rates (units of time^-1).

Let $c_A, c_{AA}, c_{AB}, c_{BB}, c_B$ denote the concentrations of the five molecular species.

Because the concentration of $A$ is constant:
**Equation 1:** $s_A + 2r_1 c_{AA} + r_5 c_{AB} = d_A c_A + 2r_2 c_A^2 + r_6 c_A c_B$

Because the concentration of $AA$ is constant:
**Equation 2:** $r_2 c_A^2 = d_{AA} c_{AA} + r_1 c_{AA}$

Because the concentration of $AB$ is constant:
**Equation 3:** $r_6 c_A c_B = d_{AB} c_{AB} + r_5 c_{AB}$

**A/B symmetry**: The reaction network is symmetric with respect to the two lists of variables $[c_A, c_{AA}, s_A, d_A, r_1, r_2, d_{AA}]$ and $[c_B, c_{BB}, s_B, d_B, r_3, r_4, d_{BB}]$ . This means that any equation will remain true if we simultaneously replace each of these parameters by the corresponding parameter from the other list.

Starting from equation 1, we can get an expression for $c_{AA}$ :

$$s_A + 2r_1 c_{AA} + r_5 c_{AB} = d_A c_A + 2r_2 c_A^2 + r_6 c_A c_B$$

$$c_{AA} = \frac{d_A c_A + 2r_2 c_A^2 + r_6 c_A c_B - s_A - r_5 c_{AB}}{2r_1}$$

Starting from equation 2, we can get another expression for $c_{AA}$ :

$$r_2 c_A^2 = d_{AA} c_{AA} + r_1 c_{AA}$$

$$c_{AA} = \frac{r_2}{d_{AA}+r_1} c_A^2$$

With $k_{AA}$ defined as $\frac{r_2}{d_{AA}+r_1}$ :

$$c_{AA} = k_{AA} c_A^2$$

By equating these two expressions of $c_{AA}$ , we can get an expression for $c_{AB}$ :

$$k_{AA} c_A^2 = \frac{d_A c_A + 2r_2 c_A^2 + r_6 c_A c_B - s_A - r_5 c_{AB}}{2r_1}$$

$$2r_1 k_{AA} c_A^2 = d_A c_A + 2r_2 c_A^2 + r_6 c_A c_B - s_A - r_5 c_{AB}$$

$$r_5 c_{AB} = d_A c_A + 2r_2 c_A^2 + r_6 c_A c_B - s_A - 2r_1 k_{AA} c_A^2$$

$$r_5 c_{AB} = d_A c_A + 2(r_2 - r_1 k_{AA}) c_A^2 + r_6 c_A c_B - s_A$$

$$c_{AB} = \frac{d_A c_A + 2(r_2 - r_1 k_{AA}) c_A^2 + r_6 c_A c_B - s_A}{r_5}$$

From the definition $k_{AA} = \frac{r_2}{d_{AA}+r_1}$ , we get $r_2 - r_1 k_{AA} = d_{AA} k_{AA}$ and therefore:

$$c_{AB} = \frac{d_A c_A + 2d_{AA} k_{AA} c_A^2 + r_6 c_A c_B - s_A}{r_5}$$

Starting from equation 3, we can get another expression for $c_{AB}$ :

$$r_6 c_A c_B = d_{AB} c_{AB} + r_5 c_{AB}$$

$$c_{AB} = \frac{r_6}{d_{AB}+r_5} c_A c_B$$

With $k_{AB}$ defined as $\frac{r_6}{d_{AB}+r_5}$ :

$$c_{AB} = k_{AB} c_A c_B$$

By equating these two values of $c_{AB}$ , we can get an expression for $c_B$ :

$$k_{AB} c_A c_B \;=\; \frac{d_A c_A \;+\; 2 d_{AA} k_{AA} c_A^2 \;+\; r_6 c_A c_B \;-\; s_A}{r_5}$$

$$r_5 k_{AB} c_A c_B \;=\; d_A c_A \;+\; 2 d_{AA} k_{AA} c_A^2 \;+\; r_6 c_A c_B \;-\; s_A$$

$$(r_5 k_{AB} - r_6) c_A c_B \;=\; d_A c_A \;+\; 2 d_{AA} k_{AA} c_A^2 \;-\; s_A$$

$$c_B \;=\; \frac{d_A c_A \;+\; 2 d_{AA} k_{AA} c_A^2 \;-\; s_A}{(r_5 k_{AB} - r_6) c_A}$$

From the definition $k_{AA} \;=\; \frac{r_6}{d_{AB} + r_5}$ , we get $r_6 - r_5 k_{AB} \;=\; d_{AB} k_{AB}$ and therefore:

$$c_B \;=\; \frac{d_A c_A \;+\; 2 d_{AA} k_{AA} c_A^2 \;-\; s_A}{-d_{AB} k_{AB} c_A}$$

By applying the A/B symmetry to this equation, we can get another expression for $c_B$ :

$$c_A \;=\; \frac{d_B c_B \;+\; 2 d_{BB} k_{BB} c_B^2 \;-\; s_B}{-d_{AB} k_{AB} c_B}$$

where $k_{BB}$ is defined as $\frac{r_4}{d_{BB} + r_3}$

$$-d_{AB} k_{AB} c_B c_A \;=\; d_B c_B \;+\; 2 d_{BB} k_{BB} c_B^2 \;-\; s_B$$

$$0 \;=\; (d_B + d_{AB} k_{AB} c_A) c_B \;+\; 2 d_{BB} k_{BB} c_B^2 \;-\; s_B$$

$$0 \;=\; 2 d_{BB} k_{BB} c_B^2 \;+\; (d_B + d_{AB} k_{AB} c_A) c_B \;-\; s_B$$

Applying the quadratic formula:

$$c_B \;=\; \frac{-b \pm \sqrt{b^2 - 4ac}}{2a} \;=\; \frac{-(d_B + d_{AB} k_{AB} c_A) \;\pm\; \sqrt{(d_B + d_{AB} k_{AB} c_A)^2 \;+\; 8 d_{BB} k_{BB} s_B}}{4 d_{BB} k_{BB}}$$

In the formula above, the coefficients are such that $a > 0$ , $b \geq 0$ , $c \leq 0$ and thus $b \;\leq\; \sqrt{b^2 - 4ac}$ . It follows that $-b \;+\; \sqrt{b^2 - 4ac}$ is non-negative, while $-b \;-\; \sqrt{b^2 - 4ac}$ is non-positive. We therefore choose $-b \;+\; \sqrt{b^2 - 4ac}$ to obtain a non-negative value of $c_B$ .

$$c_B \;=\; \frac{-(d_B + d_{AB} k_{AB} c_A) \;+\; \sqrt{(d_B + d_{AB} k_{AB} c_A)^2 \;+\; 8 d_{BB} k_{BB} s_B}}{4 d_{BB} k_{BB}}$$

By equating the two expressions that we obtained for $c_B$ , we get:

$$\frac{d_A c_A + 2 d_{AA} k_{AA} c_A^2 - s_A}{-d_{AB} k_{AB} c_A} = \frac{-(d_B + d_{AB} k_{AB} c_A) + \sqrt{(d_B + d_{AB} k_{AB} c_A)^2 + 8 d_{BB} k_{BB} s_B}}{4 d_{BB} k_{BB}}$$

$$\frac{s_A - d_A c_A - 2 d_{AA} k_{AA} c_A^2}{d_{AB} k_{AB} c_A} = \frac{-(d_B + d_{AB} k_{AB} c_A) + \sqrt{(d_B + d_{AB} k_{AB} c_A)^2 + 8 d_{BB} k_{BB} s_B}}{4 d_{BB} k_{BB}}$$

Taking $x = d_{AA} k_{AA}$ , $y = d_{BB} k_{BB}$ and $z = d_{AB} k_{AB}$ , we obtain:

$$\frac{s_A - d_A c_A - 2 x c_A^2}{z c_A} = \frac{-(d_B + z c_A) + \sqrt{(d_B + z c_A)^2 + 8 y s_B}}{4 y}$$

$$4 y s_A - 4 y d_A c_A - 8 y x c_A^2 = -d_B z c_A - z^2 c_A^2 + \sqrt{(d_B z c_A + z^2 c_A^2)^2 + 8 y s_B z^2 c_A^2}$$

$$(d_B z c_A + z^2 c_A^2) + (4 y s_A - 4 y d_A c_A - 8 y x c_A^2) = \sqrt{(d_B z c_A + z^2 c_A^2)^2 + 8 y s_B z^2 c_A^2}$$

$$\left((d_B z c_A + z^2 c_A^2) + (4 y s_A - 4 y d_A c_A - 8 y x c_A^2)\right)^2 = (d_B z c_A + z^2 c_A^2)^2 + 8 y s_B z^2 c_A^2$$

$$2(d_B z c_A + z^2 c_A^2)(4 y s_A - 4 y d_A c_A - 8 y x c_A^2) + (4 y s_A - 4 y d_A c_A - 8 y x c_A^2)^2 = 8 y s_B z^2 c_A^2$$

$$8d_B s_A y z c_A - 8d_A d_B y z c_A^2 - 16 d_B x y z c_A^3 + 8 s_A y z^2 c_A^2 - 8 d_A y z^2 c_A^3 - 16 x y z^2 c_A^4$$

$$+$$

$$16 s_A^2 y^2 - 16 s_A d_A y^2 c_A - 32 s_A x y^2 c_A^2 - 16 s_A d_A y^2 c_A + 16 d_A^2 y^2 c_A^2 + 32 d_A x y^2 c_A^3$$

$$- 32 s_A x y^2 c_A^2 + 32 d_A x y^2 c_A^3 + 64 x^2 y^2 c_A^4$$

$$=$$

$$8 y s_B z^2 c_A^2$$

$$8d_B s_A y z c_A - 8d_A d_B y z c_A^2 - 16 d_B x y z c_A^3 + 8 s_A y z^2 c_A^2 - 8 d_A y z^2 c_A^3 - 16 x y z^2 c_A^4$$

$$+$$

$$16 s_A^2 y^2 - 16 s_A d_A y^2 c_A - 32 s_A x y^2 c_A^2 - 16 s_A d_A y^2 c_A + 16 d_A^2 y^2 c_A^2 + 32 d_A x y^2 c_A^3$$

$$- 32 s_A x y^2 c_A^2 + 32 d_A x y^2 c_A^3 + 64 x^2 y^2 c_A^4 - 8 y s_B z^2 c_A^2$$

$$=$$

$$0$$

Dividing by $y$ :

$$8 d_B s_A z c_A - 8 d_A d_B z c_A^2 - 16 d_B x z c_A^3 + 8 s_A z^2 c_A^2 - 8 d_A z^2 c_A^3 - 16 x z^2 c_A^4$$

$$+$$

$$16 s_A^2 y - 16 s_A d_A y c_A - 32 s_A x y c_A^2 - 16 s_A d_A y c_A + 16 d_A^2 y c_A^2 + 32 d_A x y c_A^3$$

$$- 32 s_A x y c_A^2 + 32 d_A x y c_A^3 + 64 x^2 y c_A^4 - 8 s_B z^2 c_A^2$$

$$=$$

$$0$$

Dividing by 8:

$$d_B s_A z c_A - d_A d_B z c_A^2 - 2 d_B x z c_A^3 + s_A z^2 c_A^2 - d_A z^2 c_A^3 - 2 x z^2 c_A^4$$

$$+$$

$$2 s_A^2 y - 2 s_A d_A y c_A - 4 s_A x y c_A^2 - 2 s_A d_A y c_A + 2 d_A^2 y c_A^2 + 4 d_A x y c_A^3$$

$$- 4 s_A x y c_A^2 + 4 d_A x y c_A^3 + 8 x^2 y c_A^4 - s_B z^2 c_A^2$$

$$=$$

$$0$$

$$0 = (8x^2 y - 2xz^2)c_A^4 + (8d_A xy - 2d_B xz - d_A z^2)c_A^3$$
$$+ (s_A z^2 - d_A d_B z - 8s_A xy + 2d_A^2 y - s_B z^2)c_A^2 + (s_A d_B z - 4s_A d_A y)c_A + 2s_A^2 y$$

$$0 = 2x(4xy - z^2)c_A^4 + (d_A(8xy - z^2) - 2d_B xz)c_A^3$$
$$+ (2yd_A^2 + s_A(z^2 - 8xy) - d_A d_B z - z^2 s_B)c_A^2 + s_A(d_B z - 4d_A y)c_A + 2s_A^2 y$$

Dividing by $z$ :

$$0 = 2x\left(4\frac{xy}{z} - z\right)c_A^4 + \left(d_A\left(8\frac{xy}{z} - z\right) - 2d_B x\right)c_A^3$$
$$+ \left(\frac{2yd_A^2}{z} + s_A\left(z - 8\frac{xy}{z}\right) - d_A d_B - zs_B\right)c_A^2 + s_A\left(d_B - 4d_A\frac{y}{z}\right)c_A + 2s_A^2\frac{y}{z}$$

Eliminating $x$ , $y$ and $z$ using their definitions $x = d_{AA}k_{AA}$ , $y = d_{BB}k_{BB}$ and $z = d_{AB}k_{AB}$ , we obtain:

$$0$$
$$=$$
$$2d_{AA}k_{AA}\left(4\frac{d_{AA}k_{AA}d_{BB}k_{BB}}{d_{AB}k_{AB}} - d_{AB}k_{AB}\right)c_A^4$$
$$+$$
$$\left(d_A\left(8\frac{d_{AA}k_{AA}d_{BB}k_{BB}}{d_{AB}k_{AB}} - d_{AB}k_{AB}\right) - 2d_B d_{AA}k_{AA}\right)c_A^3$$
$$+$$
$$\left(\frac{2d_{BB}k_{BB}d_A^2}{d_{AB}k_{AB}} + s_A\left(d_{AB}k_{AB} - 8\frac{d_{AA}k_{AA}d_{BB}k_{BB}}{d_{AB}k_{AB}}\right) - d_A d_B - d_{AB}k_{AB}s_B\right)c_A^2$$
$$+$$
$$s_A\left(d_B - 4d_A\frac{d_{BB}k_{BB}}{d_{AB}k_{AB}}\right)c_A$$
$$+$$
$$2s_A^2\frac{d_{BB}k_{BB}}{d_{AB}k_{AB}}$$

This formula can be used to compute the values of the coefficients of the 4th degree polynomial given the values of the chemical parameters. The resulting values of the coefficients can then be used by a numerical solver to compute the 4 possible values of $c_A$ (the 4 roots of the polynomial). The corresponding steady-state concentrations of the other molecular species can then be computed using the formulas derived above.

$$c_B = \frac{d_A c_A + 2 d_{AA} k_{AA} c_A^2 - s_A}{-d_{AB} k_{AB} c_A} = \frac{s_A - 2 d_{AA} k_{AA} c_A^2 - d_A c_A}{d_{AB} k_{AB} c_A} = \frac{s_A / c_A - 2 d_{AA} k_{AA} c_A - d_A}{d_{AB} k_{AB}}$$

$$c_{AA} = k_{AA} c_A^2 = \frac{r_2}{d_{AA} + r_1} c_A^2$$

$$c_{BB} = k_{BB} c_B^2 = \frac{r_4}{d_{BB} + r_3} c_B^2$$

$$c_{AB} = k_{AB} c_A c_B = \frac{r_6}{d_{AB} + r_5} c_A c_B$$

Among the 4 possible values of $c_A$, the one that results in positive values for the concentrations of all five molecular species is the physically correct solution.
